# γ-secretase facilitates retromer-mediated retrograde transport

**DOI:** 10.1101/2024.06.07.597932

**Authors:** Yuka Takeo, Mac Crite, Daniel DiMaio

## Abstract

The retromer complex mediates retrograde transport of protein cargos from endosomes to the trans-Golgi network (TGN). γ-secretase is a multisubunit protease that cleaves the transmembrane domain of its target proteins. Mutations in genes encoding subunits of retromer or γ-secretase can cause familial Alzheimer disease (AD) and other degenerative neurological diseases. It has been reported that retromer interacts with γ-secretase, but the consequences of this interaction are not known. Here, we report that retromer-mediated retrograde protein trafficking in cultured human epithelial cells is impaired by inhibition of γ-secretase activity or by genetic elimination of γ-secretase. γ-secretase inhibitor XXI and knockout of PS1, the catalytic subunit of γ-secretase, inhibit endosome to TGN trafficking of retromer-dependent retrograde cargos, divalent metal transporter 1 isoform II (DMT1-II), cation-independent mannose-6-phosphate receptor (CIMPR), and shiga toxin. Trafficking of retromer-independent cargos, such as cholera toxin and a CIMPR mutant that does not bind to retromer was not affected by γ-secretase inhibition. XXI treatment and PS1 KO inhibit interaction of γ-secretase with retromer but do not inhibit the association of cargo with retromer or with γ-secretase in intact cells. Similarly, these treatments do not affect the level of Rab7-GTP, which regulates retromer-cargo interaction. These results suggest that the γ-secretase-retromer interaction facilitates retromer-mediated retrograde trafficking.

## Introduction

Retromer is a highly conserved cytoplasmic protein complex that mediates intracellular trafficking of certain transmembrane (TM) proteins, ensuring their proper localization, sorting and recycling. Retromer regulates both retrograde transport of cargo from endosomes to the *trans*-Golgi network (TGN), as well as anterograde transport from endosomes to the plasma membrane [1-3]. The absence of normal retromer function can lead to cellular dysfunction and contribute to the development of neurodegenerative diseases such as Alzheimer disease (AD) and Parkinson disease (PD) [4-8].

In mammals retromer is composed of two major subcomplexes, a core cargo recognition trimer of vacuolar protein sorting-associated proteins 26, 29, and 35 (VPS26, VPS29, and VPS35) that binds directly to protein cargo and a membrane-associated sorting nexin (SNX) dimer [2, 9]. The large family of SNXs provide diversity in terms of cargo recognition and downstream trafficking pathways. Three distinct forms of retromer complex are currently recognized: SNX-Bin, Amphiphysin, and Rvs-retromer (SNX-BAR-retromer); SNX3-retromer; and SNX27-retromer [10, 11]. Rab7 is a small GTPase that regulates intracellular protein trafficking by cycling between a membrane-associated guanosine triphosphate (GTP)-bound form and a cytosolic guanosine diphosphate (GDP)-bound form [12, 13]. When bound to GTP, Rab7 recruits the retromer complex to the endosome membrane with its TM cargo and SNX proteins [14, 15]. The retromer-SNX complex induces tubulation of the endosome membrane, leading to the formation of a vesicle containing cargo that is transported to the TGN where membrane fusion delivers the cargo to the TGN membrane [16]. The GTPase activating protein (GAP), TBC1D5, stimulates generation of GDP-Rab7 and the dissociation of retromer and its cargo from the endosome membrane during trafficking [17].

Retromer cargo proteins are TM proteins that typically possess a short sorting signal in their cytoplasmic domain that binds directly with the retromer for proper transport [18]. Cation-independent mannose 6-phosphate receptor (CIMPR) and divalent metal transporter 1 isoform II (DMT1-II) are well-studied cellular proteins that are transported in a retrograde direction by retromer from the endosome to the TGN. DMT1-II plays a role in the efficient and rapid uptake of iron across the endosomal membrane in the transferrin cycle. SNX3 binds retromer at the interface of VPS35 and VPS26. Upon binding to SNX3, VPS26 undergoes a conformational change in its cargo-binding motif, which allows recognition of the cytoplasmic domain of DMT1-II by VPS26 and SNX3 [19]. CIMPR associates with lysosomal enzymes and is transported from the Golgi to endosomes via vesicle transport. Subsequently, CIMPR is returned to the TGN by retrograde transport for reuse [20]. The role of retromer in the retrograde sorting of the CIMPR is complex. Although retromer can bind to a heterodimeric SNX-BAR membrane-remodeling complex to recognize and transport CIMPR, SNX-BARs can also directly bind CIMPR independently of retromer for delivery into the retrograde transport pathway [21, 22]. Shiga toxin is a bacterial protein produced by the bacterium, *Shigella dysenteriae*. Shiga toxin B-subunit (STxB) is transported in a retrograde fashion when it enters cells. Clathrin initiates retrograde sorting on early endosomes, and retromer generates vesicles that transport STxB to the TGN [23]. Transport of STxB requires a membrane-bound coat containing SNX1, one of the components of the retromer complex, for efficient retrograde transport. In contrast, transport of the B subunit of cholera toxin (CTxB) is not inhibited by SNX1 depletion, implying that it is retromer independent [24].

γ-secretase is a protease complex composed of four subunits: Presenilin 1 (PS1), nicastrin (NCT), anterior pharynx defective 1 (APH1), and presenilin enhancer 2 (PEN2), each of which contains at least one TM domain (TMD) [25-27]. γ-secretase specifically recognizes and cleaves substrate proteins such as Notch and the amyloid precursor protein (APP) within their TMD [28, 29]. In addition to its action as a protease, various non-proteolytic activities of γ-secretase have been described [30, 31]. AD is associated with the improper cleavage of APP by γ-secretase, leading to increased production of the amyloidogenic Aβ-42 peptide [32]. Numerous mutations within all four subunits of the γ-secretase complex can cause AD, with a majority of these mutations found in PS1, the catalytic subunit [33]. Retromer dysfunction has also been implicated in AD and PD pathogenesis [4, 34-36]. For example, reduced expression of both VPS35 and VPS26 has been observed in the postmortem brains of AD patients and mutations in VPS35, VPS29, or VPS26 are associated with the development of PD.

A stable complex between retromer and γ-secretase can be detected by co-immunoprecipitation from detergent extracts of neuronal cells, but the consequence of this interaction is not known [37]. Moreover, both γ-secretase and APP are recycled via retrograde trafficking, although it remains unclear whether their recycling depends on retromer [38]. In addition, both retromer and γ-secretase are required for efficient retrograde trafficking of human papillomaviruses (HPVs) during virus entry into cells [31, 39-42]. Therefore, we decided to test whether γ-secretase is required for the retromer-mediated retrograde trafficking of cellular cargo in uninfected cells.

In this study, we show that inhibition of γ-secretase activity and PS1 KO caused accumulation of three different retromer cargos in the endosome and their depletion from the TGN. The γ-secretase inhibitor inhibited the association between γ-secretase and retromer. Inhibition of trafficking does not appear to be due to repression of expression of the core retromer subunits, relocalization of retromer, inhibition of the association of the cargo with retromer, or perturbations in the level or nucleotide loading of Rab7. Our findings suggest that γ-secretase supports retrograde trafficking of cellular protein cargos by interacting with retromer and modulating retromer activity.

## Results

### γ-secretase is required for efficient retromer-mediated trafficking of cellular protein cargos

To determine whether γ-secretase plays a role in retromer-mediated trafficking of cellular proteins, we used immunofluorescence to monitor the effect of γ-secretase inhibition on DMT1-II trafficking. PS1 KO HeLa cells, control parental HeLa AAVS cells, and AAVS cells treated with γ-secretase inhibitor, XXI, were transfected with plasmids expressing GFP-tagged DMT1-II. As assessed by immunofluorescence and confocal microscopy at 24 hours post transfection (hpt), XXI treatment and PS1 knockout had no obvious effect on localization of EEA1 and TGN46, markers of the early endosome the TGN, respectively (Fig. 1). Notably, XXI treatment and PS1 KO increased the co-localization of GFP-DMT1-II and EEA1 compared to control cells (Fig. 1A) and decreased co-localization of GFP-DMT1-II and TGN46 (Fig. 1B), implying that γ-secretase is required for optimal retrograde trafficking of DMT1-II.

**Figure 1.**
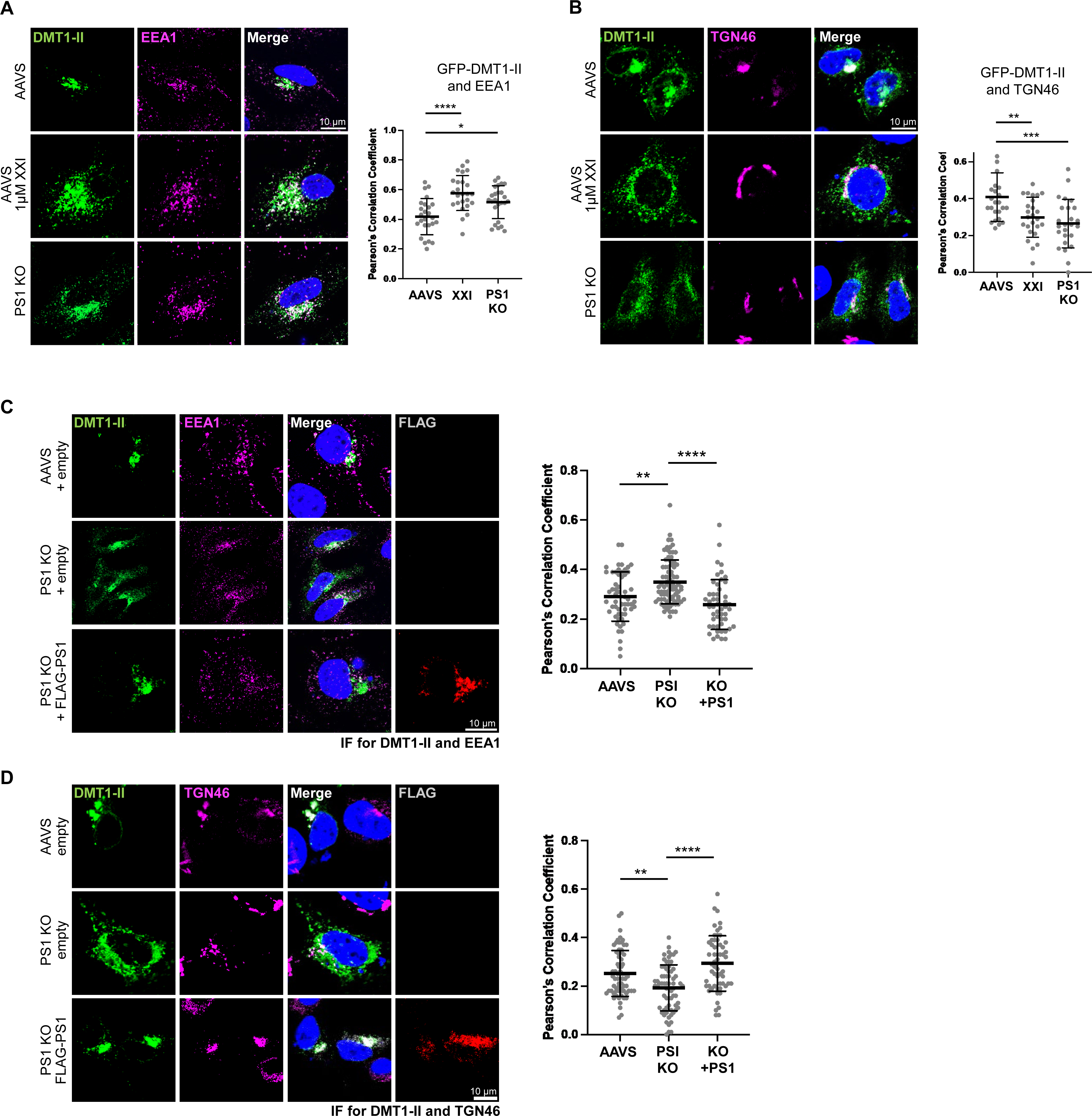
γ-secretase inhibition and PS1 knockout inhibit retrograde trafficking of DMT1-II. (**A**) Parental control HeLa cells (AAVS) and PS1 knockout HeLa cells (PS1 KO) were treated with DMSO or 1μM γ-secretase inhibitor XXI for 30 min and then transfected with a plasmid expressing GFP-DMT1-II. Cells were fixed 24 h post transfection (hpt) and stained with DAPI and an antibody recognizing EEA1. Fluorescence images of single confocal planes are shown: GFP-DMT1-II, intrinsic green fluorescence; EEA1, magenta; nuclei, blue. Merged images show overlap between GFP-DMT1-II and EEA1 pseudocolored white. Graph shows Pearson’s correlation coefficients for colocalization of GFP-DMT1-II and EEA1 in cells expressing GFP-DMT1-II. Each dot represents an individual cell (n=25), and black horizonal line indicates the mean value of the analyzed population in each group. *, *P*<0.05; ****, *P*<0.0001. Similar results were obtained in three independent experiments. (**B**) As in (**A**) except cells were stained with an antibody recognizing TGN46 instead of EEA1, and merged image shows overlap between GFP-DMT1-II and TNG46. **, *P*<0.01; ***, *P*<0.001. (**C**) AAVS and PS1 KO cells were transfected with a plasmid expressing GFP-DMT1-II and cotransfected with the empty control plasmid or a plasmid expressing FLAG-PS1. Cells were fixed 24 hpt and stained with DAPI and antibodies recognizing EEA1 and FLAG. Fluorescent images of single confocal planes are shown: GFP-DMT1-II, intrinsic green fluorescence; EEA1, magenta; FLAG-PS1, red; nuclei, blue. Merged images show overlap between GFP-DMT1-II and EEA1 pseudocolored white. Graph shows Pearson’s correlation coefficients for colocalization of GFP-DMT1-II and EEA1 in cells expressing GFP-DMT1-II (or co-expressing GFP-DMT1-II and FLAG-PS1 in the case of cells transfected with plasmid expressing FLAG-PS1 [KO + PS1]). Each dot represents an individual cell (n>50), and horizonal line indicates the mean value of the analyzed population in each group. **, *P*<0.01; ****, *P*<0.0001. Similar results were obtained in two independent experiments. (**D**) As in (**C**) except cells were stained with antibodies recognizing TGN46 and FLAG, and merged images show overlap between GFP-DMT1-II and TGN46.

We performed rescue experiments to establish that the trafficking defect in PS1 KO cells was not due to an off-target effect. For these experiments, we used PS1 fused to FLAG to allow us to identify cells expressing the introduced PS1 gene. AAVS and PS1 KO cells were transfected with the plasmid expressing GFP-DMT1-II and cotransfected with the empty control plasmid or a plasmid expressing FLAG-PS1. As expected from the results described above, in PS1 KO cells transfected with the control plasmid, GFP-DMT1-II accumulated in the EEA1 compartment at 24 hpi and was depleted from the TGN46 compartment compared to control AAVS cells (Fig. 1C and 1D). In contrast, trafficking of GFP-DMT1-II from the endosome to the TGN was restored in PS1 KO cells expressing FLAG-PS1 (Fig. 1C and 1D). These results demonstrate that the defect of DMT1-II trafficking in the PS1 KO cells is due to loss of PS1 and not due to unintended off-target effects.

To explore whether γ−secretase played a role in retromer-mediated trafficking of a second cellular protein, we examined CIMPR. As noted in the introduction, CIMPR can be transported by both retromer-dependent and retromer-independent pathways. Therefore, we first determined whether retromer knockout (KO) had a detectable effect in trafficking of CIMPR in our cells. VPS35 and VPS26 double knockout (KO) HeLa S3 (VPS35/26 KO) cells and parental control HeLa S3 cells were transduced with retrovirus expressing CD8-CIMPR, which consists of the extracellular domain of CD8 fused to the TM and cytoplasmic domain of CIMPR. Localization of CD8-CIMPR was assessed by immunofluorescence staining for CD8 and cell markers followed by confocal microscopy. Compared to control HeLa S3 cells, co-localization of CIMPR with EEA1 was increased in VPS35/26 KO cells and decreased with TGN46 (Fig. S1A and S1B), consistent with impaired endosome-to-TGN trafficking of CIMPR in the retromer KO cells.

To assess the role of γ-secretase in CIMPR trafficking, PS1 KO HeLa cells, control parental HeLa AAVS cells, and AAVS cells treated with XXI were transfected with a plasmid expressing CD8-CIMPR. γ-secretase inhibition or knockout increased CD8-CIMPR localization with EEA1 and decreased localization with TGN46 (Fig. S2A and S2B). In a rescue experiment performed as above with CD8-CIMPR, in PS1 KO cells transfected with the control plasmid, CD8-CIMPR accumulated in the EEA1 compartment at 24 hpi and was depleted from the TGN46 compartment (Fig. S2C and S2D). However, trafficking of CD8-CIMPR from the endosome to the TGN was largely restored in PS1 KO cells expressing FLAG-PS1 (Fig. S2C and S2D). Thus, as was the case for DMT1-II, the defect in retrograde trafficking of CIMPR in the PS1 KO cells is due to loss of PS1 and not due to unintended off-target effects.

The cytoplasmic domain of CIMPR present in CD8-CIMPR contains the WLM retromer binding site that mediates retromer-dependent trafficking of CIMPR from the endosome to the TGN. Substitution of the WLM sequence with AAA inhibits retrograde trafficking [20]. We determined whether trafficking of the CIMPR AAA mutant is affected by γ-secretase. AAVS, XXI-treated AAVS, and PS1 KO cells were transfected with a plasmid expressing wild-type CD8-CIMPR or the CD8-CIMPR AAA mutant. As expected, the CD8-CIMPR AAA mutant accumulated in the endosome in control cells (Fig. S1C), similar to wild-type CD8-CIMPR in cells lacking γ-secretase. XXI and PS1 KO did not affect localization of CD8-CIMPR AAA mutant. These results indicate that γ-secretase does not further affect trafficking of CIMPR when this cargo cannot bind retromer and also suggests that the inhibition of retrograde trafficking caused by γ-secretase inhibition is not the consequence of generalized cellular dysfunction.

### γ-secretase is required for efficient retromer-mediated retrograde trafficking of shiga toxin

The localization of cargos that are expressed from transfected plasmids as in the experiments described above is determined by the balance of anterograde trafficking during synthesis and maturation and retrograde trafficking. To determine the effect of γ-secretase retrograde trafficking in isolation, we studied retrograde transport of shiga toxin and cholera toxin, which utilize the retrograde pathway for cell entry [23, 24, 43]. Importantly, these toxins can be added to cells as recombinant proteins and assayed acutely for localization without the confounding effects of new synthesis or anterograde transport.

First, we tested whether retromer was involved in trafficking of the transport subunits of shiga toxin and cholera toxin (STxB and CTxB, respectively). We treated control HeLa S3 cells and VPS26/35 KO cells with fluorescent STxB or CTxB and assayed localization of these proteins 30 min later. As shown in Fig. 2A and S3A, STxB showed a tight, perinuclear localization in control cells consistent with Golgi localization with extensive overlap with TGN46. In the VPS26/35 KO cells, STxB displayed a more diffuse distribution and was markedly depleted in TGN46 compartments compared to control cells. In contrast, the distribution of CTxB and its transport to the TGN not affected by retromer knockout (Fig. 2B and S3B). These results show that retromer is required for retrograde trafficking of STxB but not CTxB into the TGN46 compartment.

**Figure 2.**
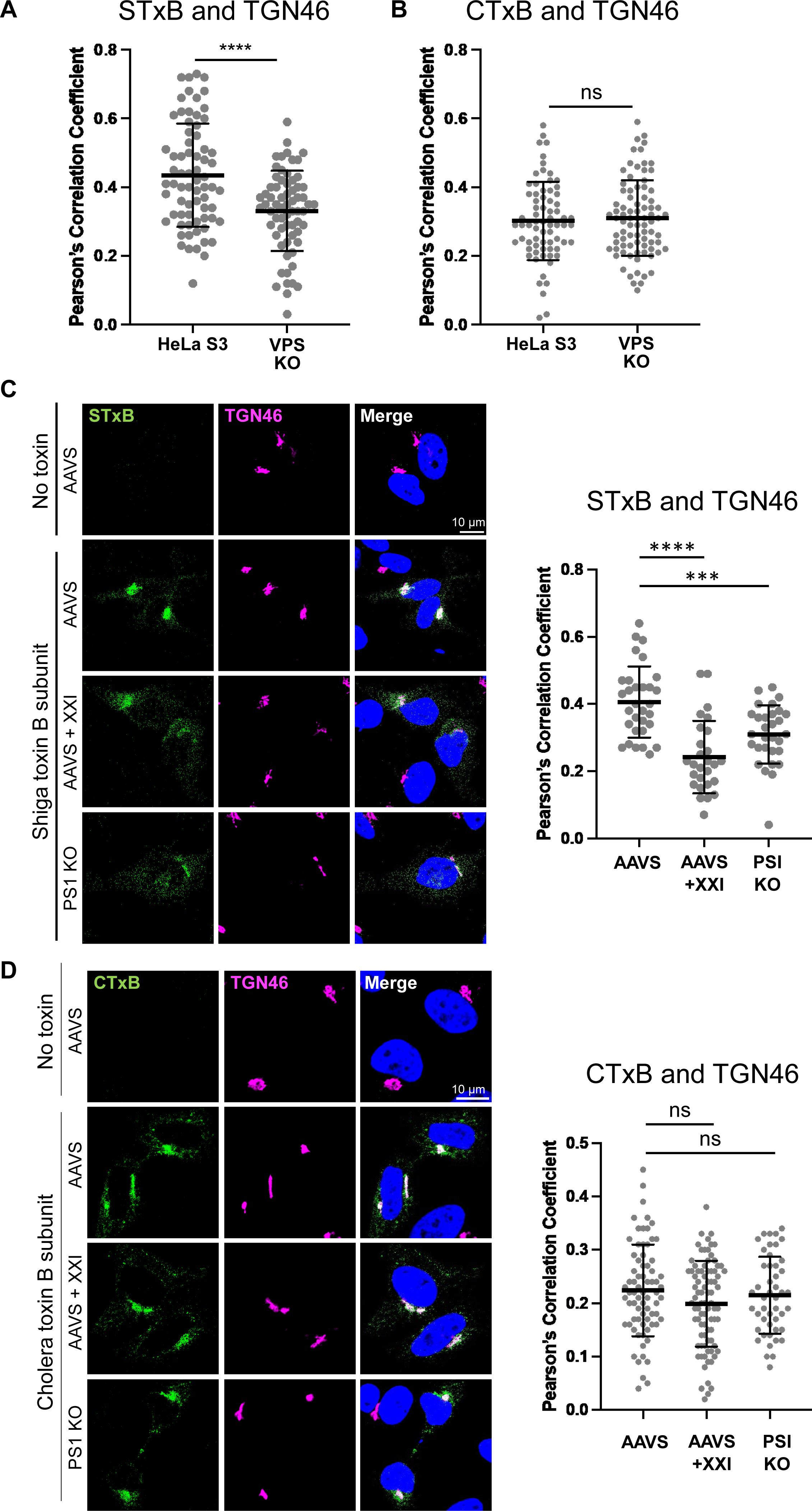
γ-secretase inhibition and PS1 knockout inhibit retrograde trafficking of shiga toxin but not cholera toxin. (**A**) HeLa S3 cells and VPS35/26 KO HeLa S3 cells (VPS KO) were incubated with 1 μg/ml fluorescent shiga toxin subunit B (STxB). Cells were fixed 30 min after the treatment and stained with DAPI and an antibody recognizing TGN46. Fluorescent confocal images were captured to determine overlap between STxB with TGN46 (representative images in Fig. S3). Graph shows Pearson’s correlation coefficients for colocalization between STxB and TGN46 in cells containing detectable toxin. Each dot represents an individual cell (n>50), and horizonal lines indicate the mean value of the analyzed population in each group. ****, P<0.0001. Similar results were obtained in two independent experiments. (**B**) Same as (A) except cells were incubated with 1 μg/ml fluorescent cholera toxin subunit B (CTxB), and Graph shows Pearson’s correlation coefficients for colocalization between CTxB and TGN46 in cells containing detectable toxin ns, not significant. (**C**) AAVS and PS1 KO HeLa cells were treated with DMSO or XXI for 24 h and then incubated with or without 1 μg/ml fluorescent STxB. Cells were fixed 30 min after treatment and stained with DAPI and an antibody recognizing TGN46. Fluorescent images of single confocal planes are shown: STxB, green; TGN46, magenta; nuclei, blue. Merged images show overlap between STxB and TGN46 pseudocolored white. Similar results were obtained in two independent experiments. Graph shows Pearson’s correlation coefficients for colocalization between STxB and TGN46 in cells containing detectable toxin. Each dot represents an individual cell (n>50), and horizonal lines indicate the mean value of the analyzed population in each group. ***, P<0.001; ****, P<0.0001. Similar results were obtained in two independent experiments. (**D**) As in (**C**) except cells were incubated with 1 μg/ml fluorescent (CTxB), and merged images show overlap between CTxB and TGN46. ns, not significant.

We next examined the acute effect of γ-secretase inhibitor on STxB and CTxB trafficking. AAVS, XXI-treated AAVS, and PS1 KO cells were incubated with STxB for 30 min, and colocalization of STxB with TGN46 was analyzed by immunofluorescence. Compared to parental cells, XXI treatment and PS1 KO caused depletion of STxB from the TGN (Fig. 2C), similar to the effect of retromer KO, indicating that γ-secretase is important for proper trafficking of STxB, a retromer-dependent retrograde cargo. In contrast, γ-secretase inhibition or knockout did not affect trafficking of CTxB, a retromer-independent cargo (Fig. 2D). These data show that γ-secretase is specifically required for efficient retromer-mediated retrograde trafficking of STxB and provide further evidence that γ-secretase inhibition does not cause a global disruption of retrograde trafficking.

We performed a rescue experiment to confirm the role of γ-secretase in trafficking of STxB. PS1 KO cells were transfected with empty vector or with the vector expressing wild-type PS1 or the PS1 L166P mutant that lacks protease activity [44, 45]. 24 h after transfection, cells were treated with fluorescent STxB for 30 min, stained with antibodies recognizing TGN46 and FLAG, and examined by confocal microscopy. As expected, in the KO cells STxB displayed diffuse distribution and relatively low colocalization with TGN46 consistent with impaired STxB trafficking, and wild-type PS1 rescued the trafficking defect. Notably, the catalytically inactive mutant did not rescue (Fig. S4A), even though it was expressed at a similar level as wild-type PS1 (Fig. S4B). The ability of wild-type but not mutant PS1 to support STxB trafficking provides additional evidence that γ-secretase is required for optimal trafficking of this cargo.

### γ-secretase and retromer do not affect each other’s expression or localization

To determine the effects of γ-secretase on expression and localization of the retromer complex and vice-versa, we performed western blotting and immunofluorescence. Lysates of AAVS, XXI-treated AAVS, and PS1 KO cells were analyzed by western blotting. We first confirmed knockout of PS1 in the KO cells and showed that XXI treatment did not affect the expression of endogenous PS1 (Fig. 3A, lanes 1-3). As we expected, PS1 KO reduced expression of another γ-secretase subunit, nicastrin (NCT) (Fig. 3A, lane 3). Importantly, there was no difference in VPS35, VPS26, and VPS29 levels between AAVS, XXI-treated AAVS, or PS1 KO cells (Fig. 3A, lanes 1-3). Similarly, KO of retromer subunits VPS35 and VPS26 reduced expression of VPS29 (as expected) but did not affect expression of γ-secretase subunits PS1 and NCT (Fig. 3A, lane 4 and 5). In addition, expression of endogenous DMT1-II was not changed by XXI treatment, PS1 KO, or VPS35/26 KO (Fig. 3A). These results show that loss of γ-secretase or retromer did not affect expression of each other or of DMT1-II.

**Figure 3.**
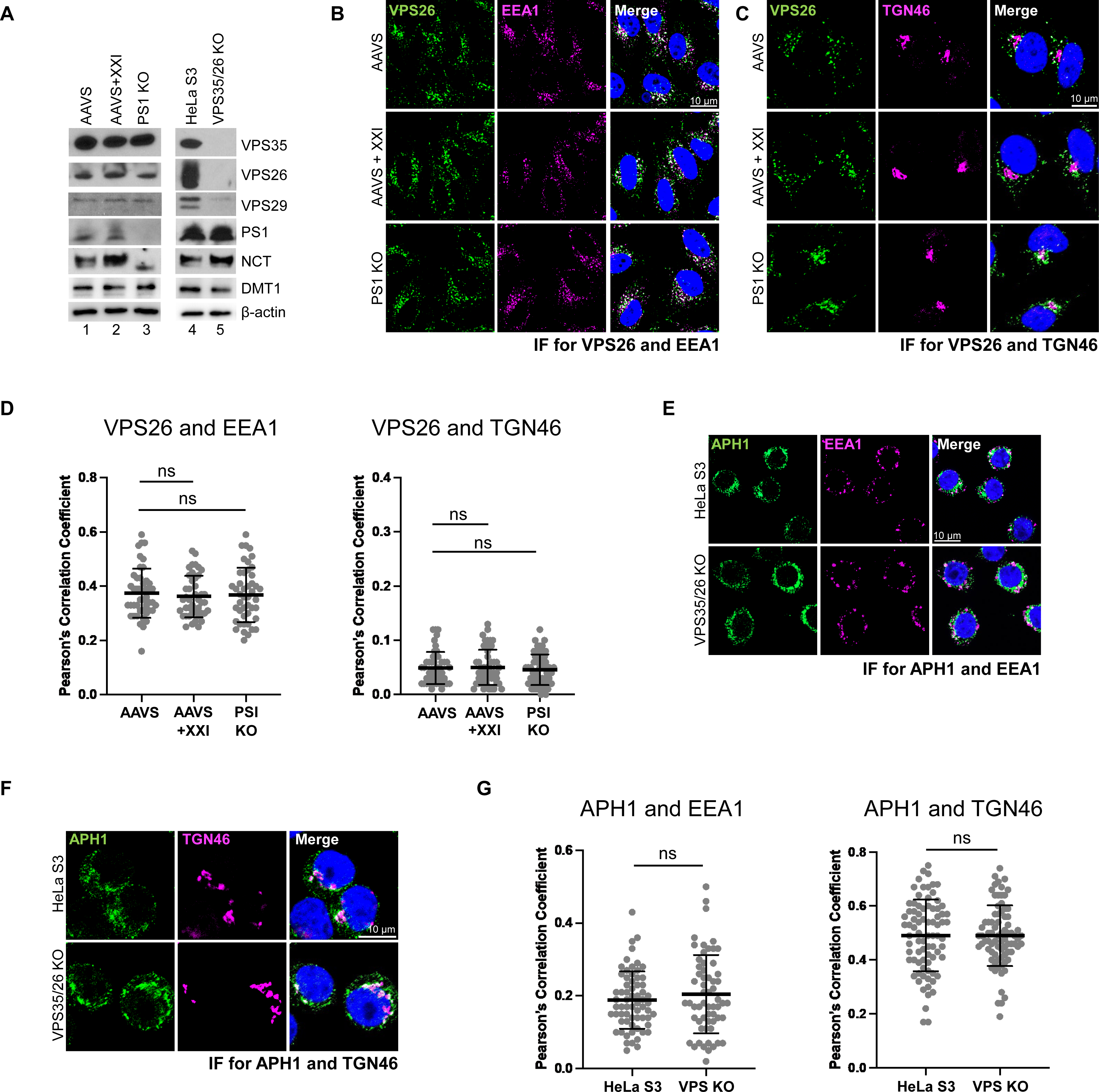
Inhibition of γ-secretase does not affect expression or localization of the retromer complex and vice-versa. (**A**) Extracts were prepared from HeLa S3, VPS35/26 KO, AAVS, and PS1 KO cells, and from AAVS cells treated with 1 μM XXI for 24 h. Cell extracts were subjected to western blot analysis using antibodies recognizing endogenous VPS35, VPS26, VPS29, PS1, NCT, DMT1, and actin (as a loading control). (**B**) AAVS and PS1 KO HeLa cells were treated with DMSO or 1 μM XXI for 24 h and then stained with DAPI and antibodies recognizing VPS26 and EEA1. Fluorescent images of single confocal planes are shown: VPS26, green; EEA1, magenta; nuclei, blue. Merged image shows overlap between VPS26 and EEA1 pseudocolored white. Graph shows Pearson’s correlation coefficients for colocalization of VPS26 and EEA1. Each dot represents an individual cell (n>40), and horizonal lines indicate the mean value of the analyzed population in each group. ns, not significant. Similar results were obtained in two independent experiments. (**C**) As in (**B**) except cells were stained with an antibody recognizing TGN46, and merged images and graph shows overlap between VPS26 and TGN46. (**D**) Images as in (**B**) and (**C**) were quantified, and graph shows Pearson’s correlation coefficient for overlap between VPS26 and EEA1 (left panel) and VPS26 and TGN46 (right panel). ns, not significant. (**E**) HeLa S3 and VPS35/26 KO cells were stained with antibodies recognizing APH1 and EEA1, and merged image shows overlap between APH1 and EEA1 pseudocolored white. (**F**) Same as (**E**) except cells were stained with antibodies recognizing APH1 and TGN46, and merged images shows overlap between APH1 and TGN46 pseudocolored white. (**G**) Images as in (**E**) and (**F**) were quantified, and graph shows Pearson’s correlation coefficient for overlap between APH1 and EEA1 (left panel) and APH1 and TGN46 (right panel). ns, not significant.

We next used immunofluorescence to monitor the localization of VPS26 and APH1 in cells. Neither XXI treatment nor PS1 KO affected co-localization of VPS26 with EEA1 or TGN46 (Fig. 3B, C and D). Similarly, VPS26/35 KO did not affect the co-localization of APH1 with EEA1 or TGN46 (Fig. 3E, F, and G). Thus, γ-secretase inhibition or KO had no apparent effect on the expression and localization of the retromer complex, and retromer KO did not affect the expression and localization of γ-secretase.

### Association of γ-secretase and retromer

There is a previous report that PS1 interacts with VPS35, as assessed by co-immunoprecipitation (co-IP) [37]. Here, we used co-IP to confirm the association of γ-secretase with retromer and determine whether γ-secretase inhibition affected the association. PS1 KO cells transfected with the empty control plasmid or a plasmid expressing FLAG-PS1were lysed in buffer containing the detergent CHAPSO and immunoprecipitated with antibody recognizing PS1. The presence of VPS35 in the immunoprecipitates was determined by western blotting. Anti-PS1 co-immunoprecipitated (co-IPed) VPS35 from PS1-transfected PS1 KO cells, but not from KO cells lacking PS1 expression, indicating that co-IP was not due to cross-reaction of the PS1 antibody with VPS35 (Fig, 4A). Next, HeLa cells treated with DMSO or XXI were lysed and immunoprecipitated with antibody recognizing PS1. Consistent with the results shown in Fig. 4A, anti-PS1 co-IPed VPS35 from DMSO-treated cells. VPS35 was also co-IPed from XXI-treated cells, but XXI reduced the association of γ-secretase with retromer (Fig. 4B). Quantitation of multiple independent repeats of this experiment showed that γ-secretase inhibition caused an approximately 75% reduction in the amount of VPS35 co-IPed.

**Figure 4.**
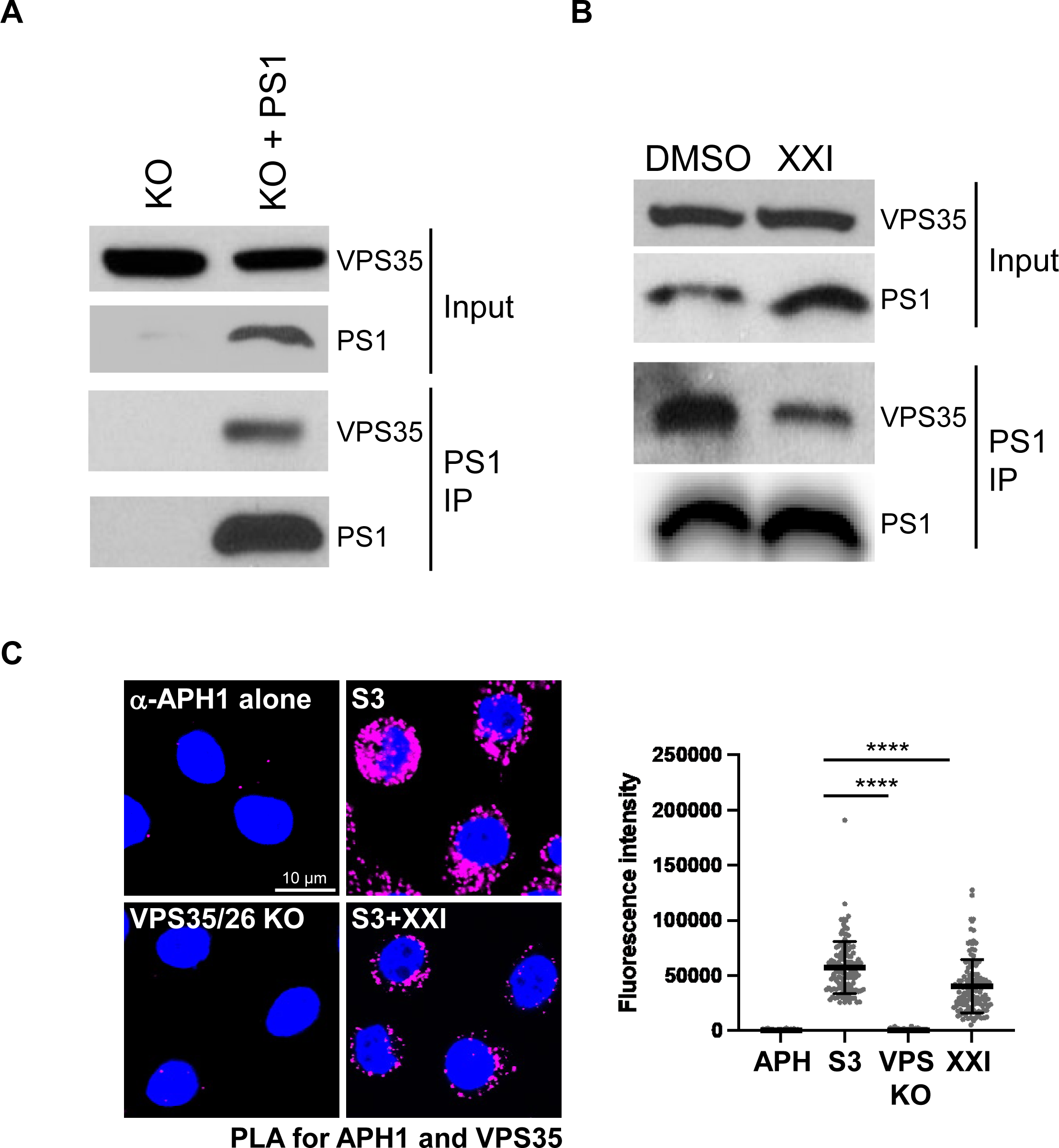
γ-secretase and retromer interact in cell lysates and intact cells. (**A**) PS1 KO cells were transfected with plasmids expressing VPS35, VPS26, and VPS29 and with empty control plasmid or a plasmid expressing FLAG-PS1. 24 hpt, detergent lysates were prepared and immunoprecipitated (IP) with an antibody recognizing PS1 and subjected to western blot analysis using an antibody recognizing VPS35 or PS1, as indicated. Blot of total lysates is shown as input. 2.25% for input and 47.75% for IP was used for Western blot analysis. Similar results were obtained in two independent experiments. (**B**) Same as (**A**) except HeLa cells expressing endogenous PS1 were transfected with plasmids expressing VPS35, VPS26, VPS29 and then treated with DMSO or 1 μM XXI. Similar results were obtained in four independent experiments. (**C**) HeLa S3 cells were treated with DMSO or 1μM XXI for 24 h, and then PLA was performed with antibodies recognizing APH1 and VPS35. As negative controls, PLA was performed in VPS26/35 KO cells or in HeLa S3 cells with the VPS35 antibody omitted (α-APH1 alone). Images show single confocal planes. PLA signals are magenta; nuclei are stained blue with DAPI. The fluorescence of PLA signals was determined from multiple images. In the graph, each dot represents an individual cell (n>50) and horizonal lines indicate the mean value of the analyzed population in each group. ****, *P*<0.0001. Similar results were obtained in two independent experiments.

To determine whether γ-secretase and retromer interacted in intact cells, we used proximity ligation assay (PLA), which generates a fluorescent signal when two proteins of interest recognized by different antibodies are within 40 nm. We performed PLA for APH1 and VPS35. As expected, only a low background level of PLA signal was detected in HeLa S3 cells incubated with APH1 antibody alone and in VPS35/26 KO cells incubated with both VPS35 and APH1 antibodies. In contrast, when antibodies recognizing both APH1 and VPS35 were used for PLA in HeLa S3 cells, clear PLA signals were observed (Fig. 4C). Although PLA cannot demonstrate a direct interaction between the proteins of interest, these data strongly suggest that γ-secretase and retromer also interact in intact cells. We also used PLA to assess whether XXI affected the interaction between γ-secretase and retromer in cells. The APH1-VPS35 PLA signal was reduced by XXI treatment (Fig. 4C). The PLA signal between VPS35 and FLAG-tagged PS1 L166P was also reduced (Fig. S4C). Thus, the co-IP and PLA results show that γ-secretase and retromer interacted in cell lysates and in intact cells and that this interaction was inhibited but not abolished by inhibition of γ-secretase activity.

### γ-secretase is not required for association of retromer with cargo in intact cells

We next explored the basis for the defect in retrograde cargo trafficking caused by inhibition of γ-secretase activity and PS1 KO. The immunofluorescence and western blotting results in Fig. 3 showed that γ-secretase inhibition did not affect the localization or expression of retromer. We hypothesized that γ-secretase controls the association of retromer with the cargo, so that inhibition or loss of γ-secretase impairs the ability of retromer to transport the cargo to the TGN from the endosome. We used PLA to test whether γ-secretase was required for the association of retromer and cargo. AAVS, XXI-treated AAVS, and PS1 KO cells were mock transfected or transfected with a plasmid expressing GFP-DMT1-II. Expression of GFP-DMT1-II was not changed by XXI or PS1 KO (Fig. 5A). PLA was then performed with antibodies recognizing GFP and VPS35. As expected, a low background level of PLA signal was detected in control AAVS cells not transfected with GFP-DMT1-II, and clear GFP-VPS35 PLA signal was detected in AAVS cells expressing GFP-DMT1-II, indicating association of VPS35 and DMT1-II (Fig. 5B). There was also abundant VPS35/GFP-DMT1-II PLA signal in XXI-treated and PS1 KO cells compared to control AAVS cells, indicating that γ-secretase was not required for the association of retromer and DMT1-II. Similarly, γ-secretase inhibition or KO did not inhibit the VPS35/CIMPR PLA signal (Fig. 5C). In fact, γ-secretase inhibition or KO resulted in increased PLA signal between retromer and both cargo proteins. These data show that inhibition of γ-secretase activity and PS1 KO do not reduce the association of the cargo and retromer.

**Figure 5.**
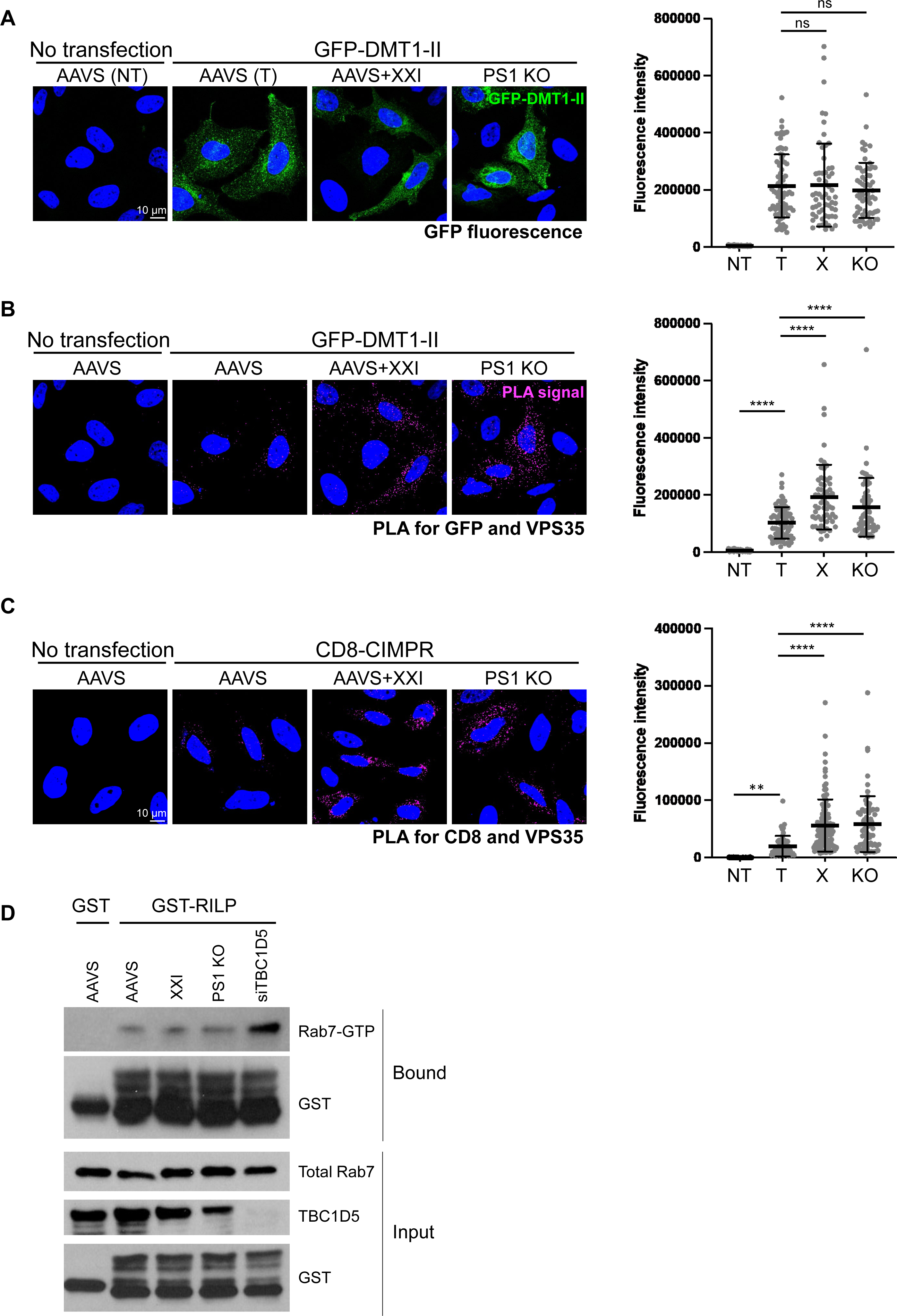
γ-secretase inhibition and PS1 knockout do not block association of retromer and cargo nor increase the levels of GTP-Rab7. **(A**) AAVS and PS1 KO HeLa cells were treated with DMSO or 1 μM XXI for 30 min and then mock transfected (NT) or transfected with a plasmid expressing GFP-DMT1-II. Cells were fixed 24 hpt and stained with DAPI. Single confocal planes are shown: GFP-DMT1-II, intrinsic green fluorescence; nuclei, blue. The GFP signals from multiple images was determined. In the graph, each dot represents an individual cell (n>50) and horizonal lines indicate the mean value of the analyzed population in each group. ns, not significant. Similar results were obtained in two independent experiments. (**B**) Cells were treated as in panel (**A**). After fixation, PLA was performed with antibodies recognizing GFP and VPS35. Single confocal planes are shown: PLA signals, magenta; nuclei, blue. These are the same fields of cells shown in panel (**A**). PLA signals from multiple images was determined in cells expressing GFP-DMT1-II. In the graph, each dot represents an individual cell (n>50) and horizonal lines indicate the mean value of the analyzed population in each group. ****, *P*<0.0001. Similar results were obtained in two independent experiments. (**C**) As in (**B**) except cells were transfected with a plasmid expressing CD8-CIMPR, and PLA was performed with antibodies recognizing CD8 and VPS35. **, *P*<0.01; ****, *P*<0.0001. (**D**) As indicated, AAVS and PS1 KO cells were untransfected, or PS1 KO cells were transfected with siRNA targeting TBC1D5 and then treated with DMSO or 1 μM XXI for 24 h. 48 hpt, detergent lysates were incubated with GST or GST-RILP, pulled down with glutathione agarose and subjected to western blot analysis by using antibodies recognizing Rab7 or GST (panels labeled bound). Western blot of total lysates probed for Rab7, TBC1D5, and GST is shown as input. Similar results were obtained in three independent experiments.

### γ-secretase inhibition does not affect levels of Rab7 or Rab7-GTP

The association of cargo with retromer during trafficking is controlled by the small GTPase, Rab7. To initiate retromer-mediated retrograde trafficking, the retromer complex and cargo are recruited to the endosome membrane under the control of GTP-Rab7. Conversion of GTP-Rab7 to GDP-Rab7 is stimulated by TBC1D5, a Rab7 GAP. Manipulations that increase the level of GTP-Rab7, e.g., expression of a constitutively active Rab7 mutant or knockdown of TBC1D5, increase association of cargo with retromer, whereas GTP hydrolysis promotes the release of retromer from cargo and endosomes [17, 46].

Because Rab7 controls the association and dissociation of cargo with retromer, we assessed the effect of γ-secretase inhibition on Rab7 levels. XXI treatment and PS1 KO did not change the abundance of total Rab7, as assessed by western blotting (Fig. 5D). To test whether γ-secretase inhibition increased the abundance of GTP-bound Rab7, which might account for the increased PLA signal between VPS35 and cargo in the inhibited cells, we performed pull-downs with recombinant Rab-interacting lysosomal protein (RILP), which binds GTP-bound but not GDP-bound Rab7. AAVS, XXI-treated AAVS, and PS1 KO cells were transfected with siRNA control, and AAVS cells were transfected with siRNA targeting the Rab7 GAP TBC1D5. Purified glutathione S-transferase (GST) or a GST-RILP fusion protein was incubated with lysates prepared from cells 48 hours after siRNA transfection. Protein complexes were pulled down with glutathione beads and blotted with anti-Rab7 antibody to detect GTP-Rab7 (Fig. 5D). As expected, in lysates from AAVS cells, GTP-Rab7 was pulled-down with GST-RILP but not with the negative control reagent GST. Knockdown of TBC1D5 caused an increase in GTP-Rab7, also as expected (compare lane 5 to lane 2). However, neither XXI treatment nor PS1 KO changed the amount of Rab7 pulled down by GST-RILP (compare lane 3 and 4 to lane 2). These results indicates that loss of γ-secretase activity did not affect the level of GTP-Rab7. Thus, changes in the level of GTP-Rab7 are not responsible for the decrease in retromer-mediated trafficking when γ-secretase is inhibited.

### γ-secretase colocalizes with STxB in intact cells

Finally, we tested whether γ-secretase and the retromer cargo STxB colocalize. We treated control HeLa S3, XXI-treated HeLa S3, and VPS35/26 KO cells with fluorescent STxB for 30 min, and localization of APH1 and STxB was analyzed by immunostaining and confocal microscopy. As shown in Fig. 6, STxB colocalized with APH1 in untreated cells. In cells treated with XXI or lacking retromer, STxB displayed more diffuse distribution, as noted in Fig. 2C and S3A, but these treatments did not disrupt the association between STxB and γ-secretase. In fact, colocalization of APH1 and STxB was increased in the retromer KO cells and the XXI-treated cells compared to control cells. These results show that γ-secretase and the retromer cargo STxB display overlapping localization in cells, which is not decreased by inhibition of γ-secretase activity or retromer KO.

**Figure 6.**
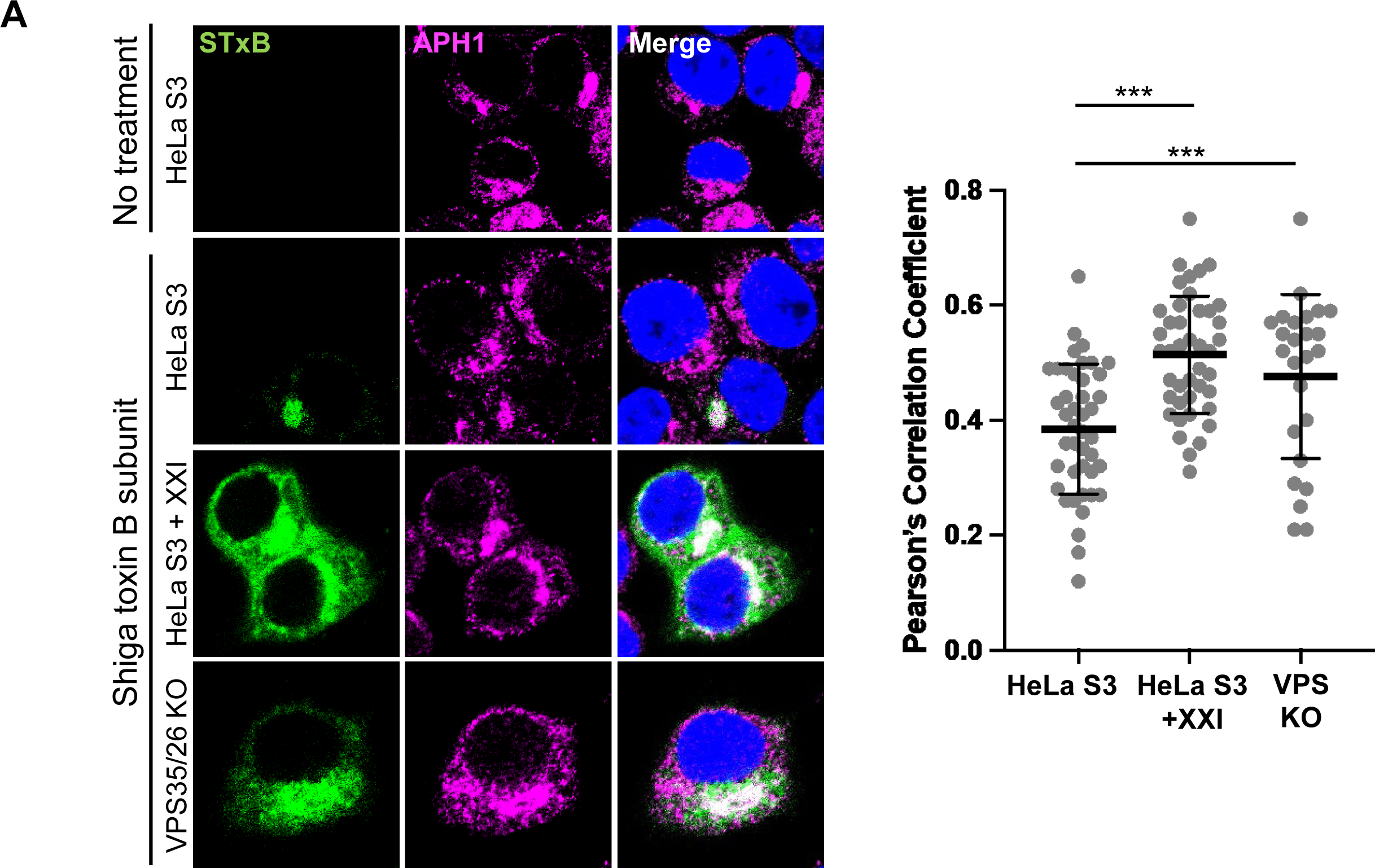
γ-secretase colocalizes with STxB in intact cells. HeLa S3 and VPS26/35 KO cells were treated with DMSO or 1 μM XXI for 24 h and then incubated with 1 μg/ml fluorescent STxB for 30 min. Cells were fixed and stained with DAPI and an antibody recognizing APH1. Fluorescent confocal images were captured to determine overlap between STxB with APH1. Graph shows Pearson’s correlation coefficients for colocalization between STxB and APH1 in cells containing detectable toxin. Each dot represents an individual cell (n>50), and horizonal lines indicate the mean value of the analyzed population in each group. ***, *P*<0.001. Similar results were obtained in two independent experiments.

## Discussion

Trafficking of intracellular proteins is under tight control to ensure proteins reach their proper destinations. Retromer is a key factor that initiates retrograde trafficking of cellular TM proteins from the endosome to the TGN, while γ-secretase plays no known role in retrograde trafficking of cellular proteins. We previously showed that retromer and γ-secretase are both required for efficient delivery of HPV into the retrograde trafficking pathway during virus entry [31, 41]. In addition, retromer and γ-secretase are reported to interact in neuronal cells, and both are implicated in the pathogenesis of neurodegenerative diseases [4, 33, 34, 37, 47]. Based on these findings, we decided to test whether γ-secretase affected retromer-mediated retrograde trafficking of cellular proteins.

Here, we used immunofluorescent imaging to show that endosome-to-TGN trafficking of three retromer cargoes was impaired when γ-secretase was inhibited or knocked out. There cargos are two human TM proteins, DMT1-II and CIMPR, which were newly synthesized from transfected plasmids, and STxB, a recombinant bacterial toxin subunit that was acutely added to cells. In contrast, γ-secretase inhibition did not affect trafficking of a CIMPR mutant that is not recognized by retromer, nor did it affect trafficking of CTxB, a bacterial toxin whose trafficking is not dependent on retromer [20, 24]. Thus, γ-secretase inhibition specifically inhibited retromer-mediated trafficking. Retrograde-mediated trafficking was inhibited but not abolished in the PS1 knockout cells, indicating that γ-secretase facilitates retromer-mediated trafficking but is not absolutely required for trafficking.

Two of the cargos we studied use different SNX-retromer complexes. STxB requires the SNX-BAR protein, SNX-1, whereas DMT1-II requires SNX-3, which lacks a BAR domain [19, 24]. The inhibition of both cargos by γ-secretase inhibition implies that γ-secretase stimulates multiple types of retromer-mediated retrograde trafficking events. Further experiments are required to determine the specific trafficking pathways that are facilitated by γ-secretase.

Another group used coIP to demonstrate that γ-secretase interacts with retromer in detergent lysates of cultured murine neuroblastoma neuro2a cells and in homogenized mouse brain [37]. Here, we showed that retromer and γ-secretase also associate in lysates of cultured human epithelial cells. Furthermore, our PLA studies imply that this interaction also occurs in intact cells. Thus, the association of retromer and γ-secretase occurs in a range of cell types and is not an artifact of cell lysis. The γ-secretase-retromer interaction was inhibited by XXI treatment or the L166P PS1 mutation, both of which decrease trafficking, suggesting that the association of γ-secretase with retromer is linked to its ability to support retrograde trafficking.

Inhibition of γ-secretase does not disrupt the association between cargo and retromer or γ-secretase. Indeed, γ-secretase inhibition appears to increase retromer-cargo and γ-secretase-cargo interaction. Retrograde transport of cargo to the TGN is a complex process that requires many proteins acting downstream of retromer-cargo interaction. For example, other proteins contribute to tubule formation and fission and vesicle movement to the TGN [48]. The association of retromer and cargo despite γ-secretase inhibition implies that γ-secretase acts relatively early in retrograde trafficking, after retromer-cargo association but before retromer-cargo dissociation and vesicle fission.

We have ruled out several potential mechanisms by which γ-secretase inhibition might impair retromer-mediated retrograde trafficking. PS1 knockout or inhibition did not appear to affect expression of any of the retromer subunits or localization of VPS26, suggesting that the inhibitory effect of impairing γ-secretase function is not due to altered expression or localization of retromer. Although retromer activity is required for proper expression and localization of PS1 in murine neuronal cells when the endocytic pathway was perturbed by treatment with agents such as NH_4_Cl that inhibit lysosomal function [37], no effects on PS1 were observed in the absence of these treatments, consistent with our results.

Our PLA studies indicate that γ-secretase is not required for association of retromer with its cargo, which is mediated by direct binding to short sequence motifs in the cargo. Similarly, γ-secretase and the STxB colocalize in intact cells, even in cells lacking retromer. Thus, γ-secretase and retromer can each associate with cargo in the absence of the other factor. In fact, γ-secretase inhibition or KO was accompanied by increased retromer-cargo association, and γ-secretase inhibition and retromer KO increased γ-secretase-cargo association. Rab7-GTP normally recruits cargo to retromer, but the increased retromer-cargo association upon γ-secretase inhibition is not due to elevated Rab7-GTP. Increased association of cargo with retromer does not necessarily inhibit trafficking. For example, expression of constitutively active Rab7 increases CIMPR-retromer association but does not inhibit trafficking [17, 49]. Therefore, we conclude that the increased retromer-cargo association is likely to be a consequence of inhibition of trafficking, not the cause of it.

It is not clear if the proteolytic activity of γ-secretase is required for trafficking. The chemical γ-secretase inhibitor XXI and the L166P PS1 mutation that abolishes catalytic activity inhibit γ-secretase cleavage activity and interfere with trafficking, but both also inhibit the interaction between γ-secretase and retromer. None of the retromer subunits contain TMDs, the substrate for cleavage, nor do the SNX proteins or Rab7. Furthermore, γ-secretase does not cleave the TMD of the cargo itself because retrograde cargo cleavage is not commonly observed during retromer-mediated trafficking, and in our experiments inhibiting γ-secretase did not affect the level of full-length DMT1-II (Fig. 3A). Thus, if the proteolytic activity of γ-secretase is involved in retrograde trafficking, it must act indirectly by cleaving some other protein(s) involved in trafficking. Alternatively, γ-secretase may play a non-proteolytic role in trafficking, such as serving as a scaffold to assemble a protein complex important for trafficking. The presence of TMDs in both the retromer cargos and γ-secretase substrates raises the possibility that any such a scaffolding function may involve interactions between TMDs.

Our findings that retromer and γ-secretase are both required for efficient retrograde trafficking may have relevance to neurodegenerative diseases. Mutations in PS1 and other genes encoding γ-secretase subunits can cause familial AD and PD [33, 50]. γ-secretase catalyzes intramembranous proteolysis of amyloid precursor protein (APP) and generates Aβ, the accumulation of which is thought to be key in the pathogenesis of AD [51]. The biochemical role of γ-secretase in PD is unclear, but there is a report that Istradefylline, an anti-PD drug, enhances Aβ generation and γ-secretase activity [52]. It is also unclear how retromer contributes to the pathogenesis these diseases, although perturbations in endocytic trafficking have been implicated in AD [6, 38, 47, 53]. Mutations in retromer subunits including VPS35 D620N can cause PD disease, and retromer levels are reduced in the brains of patients with late-onset AD [54]. The VPS35 D620N mutation increases colocalization of CIMPR with VPS35 and impairs trafficking of CIMPR and other retromer cargos [55], similar to the pattern we observed when γ-secretase is inhibited. These results raise the possibility that γ-secretase inhibition and the VPS35 D620N mutation may have similar effects on retromer-mediated trafficking. We speculate that some γ-secretase mutations may affect retrograde trafficking of a variety of retromer cargos, which could contribute to disease pathogenesis. For example, mutations in SORL1, which affects APP recycling, can cause familial AD [56]. Since SORL1 is a retromer cargo, impairment of retromer activity may affect SORL1 function. It will be interesting to determine whether some γ-secretase mutations associated with neurodegenerative disease contribute to pathogenesis by affecting retromer-mediated retrograde trafficking.

In summary, we show that γ-secretase is required for optimal retromer-mediated retrograde trafficking of cellular proteins from the endosomes to the TGN. As well as providing new insight into the regulation of intracellular protein trafficking, the activities reported here may also have implications for the pathogenesis and possibly treatment of devastating human disease.

## Acknowledgments

We thank Billy Tsai, Kashif Mehmood, Jeongjoon Choi, Christopher Burd, and Takamasa Inoue for helpful reagents and discussions. This work was supported by a grant from the NIH to D.D. and Billy Tsai (AI150897).

## Materials and methods

### Inhibitors and antibodies

| Inhibitors | Source | Catalog no. |
| --- | --- | --- |
| XXI | Sigma | 565790 |
| Shiga toxin B subunit | Sigma | SML0562 |
| Cholera toxin B subunit | Thermo Fisher Scientific | C34775 |

| Antigen | Source | Catalog no. | Species | Application | Dilution |
| --- | --- | --- | --- | --- | --- |
| EEA1 | Cell Signaling | 2411 | Rabbit | IF | 1:100 |
| EEA1 | BD | 610457 | Mouse | IF | 1:100 |
| TGN46 | Abcam | ab50595 | Rabbit | IF | 1:200 |
| CD8 | AdipoGen | Anc153-020 | Mouse | IF, PLA | 1:400 |
| VPS35 | Abcam | ab157220 | Rabbit | PLA, WB | 1:200, 1:1000 for |
| VPS35 | Abcam | ab57632 | Mouse | PLA, WB | 1:200, 1:1000 for |
| VPS26 | Abcam | ab23892 | Rabbit | IF, WB | 1:200, 1:1000 for |
| VPS29 | Abcam | ab236796 | Rabbit | WB | 1:1000 |
| PS1 | Cell Signaling | 5643 | Rabbit | WB | 1:1000 |
| NCT | Santa Cruz | sc-376513 | Mouse | WB | 1:1000 |
| APH1 | Thermo Fisher | PA1-2010 | Rabbit | IF, PLA | 1:50 |
| DMT1 | Abcam | ab55735 | Mouse | WB | 1:1000 |
| GFP | Santa Cruz | sc-9996 | Mouse | PLA | 1:100 |
| Rab7 | Cell Signaling | 9367 | Rabbit | WB | 1:1000 |
| GST-HRP | Santa Cruz | sc-138 | Mouse | WB | 1:2000 |
| Flag | Thermo Fisher | MA1-142 | Rat | IF | 1:500 |
| TBC1D5 | Abcam | Ab203896 | Rabbit | WB | 1:1000 |
| pan actin | Cell Signaling | 4968 | Rabbit | WB | 1:2000 |

| Secondary antibody (Thermo Fisher Scientific) | Catalog# |
| --- | --- |
| Donkey anti-Mouse IgG (H+L) Highly Cross-Adsorbed Secondary Antibody, Alexa Fluor™ 647 | A31571 |
| Goat anti-Rabbit IgG (H+L) Highly Cross-Adsorbed Secondary Antibody, Alexa Fluor™ 647 | A21245 |
| Goat anti-Rat IgG (H+L) Cross-Adsorbed Secondary Antibody, Alexa Fluor™ 647 | A21247 |
| Donkey anti-Rabbit IgG (H+L) Highly Cross-Adsorbed Secondary Antibody, Alexa Fluor™ 568 | A10042 |
| Donkey anti-Mouse IgG (H+L) Highly Cross-Adsorbed Secondary Antibody, Alexa Fluor™ 488 | A21202 |
| Donkey anti-Rabbit IgG (H+L) Highly Cross-Adsorbed Secondary Antibody, Alexa Fluor™ 488 | A21206 |

### Cell culture

HeLa S3 [American Type Culture Collection (ATCC), catalog no. CCL-2.2], and HeLa (ATCC, catalog no. CCL-2) cells were obtained from ATCC. All cell lines were cultured at 37℃ and 5% CO_2_ in Dulbecco’s modified Eagle’s medium (DMEM) supplemented with 10% fetal bovine serum (FBS), L-glutamine, 100 units/mL penicillin streptomycin, and 20mM HEPES (DMEM10). Cell lines were verified by using the ATCC cell authentication service.

### Plasmid construction

Plasmid expressing 3xFlag-PS1 or L166P PS1was constructed as follows: To generate the pCMVTNT-3xFlag-linker vector for subcloning, 3xFlag and linker that contains BamHI and EcoRI sites were introduced between XhoI and BamHI sites of pCMVTNT-HA-L2-3xFLAG, a gift from Billy Tsai (University of Michigan), using In-Fusion cloning (TaKaRa Bio, 638948). During In-Fusion cloning, we deleted the original BamHI site. The PS1 or L166P PS1 gene was amplified from pCAG-PS1 or pCAG-L166P PS1 (gifts from Billy Tsai) respectively, using primers 5’-GTG GTT CTG GTG GTG GAT CCG GTA CAG AGT TAC CTG CAC CGT TGT CCT ACT TC-3’ and 5’-GCT CGA AGC GGA ATT CCT AGA TAT AAA ATT GAT GGA ATG CTA ATT G-3’. The amplified segment was introduced between BamHI and EcoRI sites of pCMVTNT-3xFlag-linker by In-Fusion cloning. Resulting plasmids were confirmed by DNA sequencing.

### Generation of KO cells

To generate VPS35 KO cells, 5’-CAC CGG GAG CAT TTG CGC TTG CGG C-3’ and 5’-AAA CGC CGC AAG CGC AAA TGC TCC C-3’ comprising the VPS35 guide RNA were annealed and inserted into the pLentiCRISPRv2 (Addgene, 49535) vector. pLentiCRISPRv2 plasmid expressing the guide RNA or the empty vector was transfected into 293T cells using Lipofectamine 2000 with pMG2.D and psPAX2 packaging plasmid. 48 hpt, the growth medium containing the lentivirus was harvested and filtered through a 0.45μm filter. HeLa S3 cells were transduced with the lentivirus by using 4μg/mL polybrene. The culture medium was replaced with fresh DMEM10 at 24 hours post infection (hpi), and 0.4μg/mL puromycin was added at 48 hpi. After selection, surviving HeLa S3 cells were diluted and seeded into 96 well plates, so that each well contained no more than a single cell. Wells were monitored for single colony growth. After cell expansion, VPS35 KO was confirmed with western blotting. To generate VPS35/26 double KO cells, 5’-CAC CGT TCA TAC CTC AAG CGG ACA T-3’ and 5’-AAA CAT GTC CGC TTG AGG TAT GAA C-3’ comprising the VPS26 guide RNA were annealed and introduced into pLentiCRISPRv2 hygro (Addgene, 98291). Lentivirus was produced as described above and transduced into VPS35 KO cells. Cells were selected with 0.25 mg/mL of hygromycin B (Thermo Fisher Scientific, 10687010), cloned, and tested by western blotting for VPS26 knockout.

PS1 KO cells and control AAVS HeLa cells were gifts from Billy Tsai and Takamasa Inoue. PS1 KO cells were generated as previously described [31]. To generate AAVS control cells, 5’-CAC CGC GGG AGA TCC TTG GGG CGG T-3’ and 5’-TTT GAC CGC CCC AAG GAT CTC CCG C-3’ comprising the guide RNA of AAVS1 were annealed and introduced into pX330 vector. The plasmid was transfected into HeLa cells.

### Immunofluorescence

2x10^5^ AAVS and PS1 KO cells or 4x10^5^ HeLa S3 and VPS35/26 KO cells were plated in 24-well plates containing glass coverslips 20 h prior to transfection. Cells were treated with DMSO or 1 μM XXI 30 min before transfection with 1 μg plasmid expressing protein of interest. 24 hpt, cells were fixed with 4% paraformaldehyde (Electron Microscopy Sciences) at room temperature (RT) for 10 min, permeabilized and blocked with 0.1% saponin (Sigma Aldrich, 47036) in DMEM10 for 10-30 min at RT. Primary antibodies were diluted in 0.1% saponin in DMEM10 and incubated with the cells overnight at 4 ℃. Alexa-Fluor-conjugated secondary antibodies were also diluted in 0.1% saponin in DMEM10 and incubated at RT for 1 h. When the primary antibody recognizing VPS26 or APH1 was used, Alexa Fluor 647-conjugated anti-TGN46 antibody was prepared using Alexa Fluor Antibody Labeling Kits (Thermo Fisher Scientific, A20186) according to the manufacturer’s instructions. Coverslips were mounted with mounting medium containing DAPI (Abcam, ab104139). Cellular fluorescence was imaged using the Zeiss LSm980 confocal microscope. Images were processed using a Zeiss Zen software version 3.1 and quantified using Image J Fiji version 1.14.0/1.54f.

### GFP-DMT1-II and CD8-CIMPR trafficking assays

Cells were plated as above and treated with DMSO or 1 μM XXI for 30 min. 1 μg of pGFP-DMT1-II (obtained from Mitsuaki Tabuchi, Kagawa University) or CD8-CIMPR plasmid (obtained from Matthew Seaman, Cambridge Institute for Medical Research) was transfected into the cells by using Trans-IT HeLaMONSTER reagent (Mirus Bio). DMSO and XXI were maintained in the culture medium during transfection. 24 hpt, cells were fixed and processed for immunofluorescence as described above.

### Toxin uptake assays

Cells plated as described above were treated with DMSO or 1μM XXI for 24 h and then incubated for 30 min with 1 μg/mL of Cholera Toxin Subunit B, Alexa Fluor 488 Conjugate (Thermo Fisher Scientific, C34775) or Shiga Toxin 1 B subunit (Sigma, SML0562), conjugated with Alexa Fluor 555 using Alexa Fluor 555 Microscale Protein Labeling Kit (Thermo Fisher Scientific, A30007) according to the manufacturer’s instructions. DMSO and XXI were maintained in the culture medium during toxin incubation. Cells were fixed and processed for immunofluorescence as described above.

### Western blot analysis

Cells were lysed using ice-cold Radioimmunoprecipitation assay (RIPA) [50 mM Tris pH7.4, 150 mM NaCl, 1% Nonidet P-40, 1% sodium deoxycholate, 0.1% sodium dodecyl sulfate, 1 mM ethylenediaminetetraacetic acid] buffer supplemented with 1x HALT protease and phosphatase inhibitor cocktail (Pierce Thermo Fisher Scientific) for 15 min at 4 ℃. After centrifugation at maximum speed for 15 min in an Eppendorf 5430R centrifuge, the supernatant was mixed with 4x Laemmli dye (Bio-rad) supplemented with 10% 2-mercaptoethanol and incubated for 5 min at 100°C. Samples were then separated by SDS-PAGE (4-12% acrylamide) (Bio-rad) and analyzed by western blotting. Secondary horseradish peroxidase (HRP)-conjugated antisera recognizing rabbit or mouse antibodies as appropriate (Jackson ImmunoResearch, 711-035-152, 115-035-146) were used at 1:5000 to 1:10000 dilution. The blots were developed with SuperSignal West Pico or Femto Chemiluminescent substrate (Pierce) and visualized by using FluorChem Imager (Bio-technne, FE0685) or film processor (Fujifilm).

### Co-immunoprecipitation

3x10^6^ HeLa cells or PS1 KO HeLa cells per dish were plated in 60 mm dishes. Polyethylenimine (PEI) was used to transfect PS1 KO cells with three plasmids (3 μg each) encoding retromer subunits (VPS35, VPS26, and VPS29), originally constructed by C. Haft (NIDDK), were obtained from Jae Jung (University of Southern California) and Nam-Hyuk Cho (Seoul National University) [57], and 3 μg of a control plasmid or a plasmid expressing FLAG-tagged PS1, constructed as described above. HeLa cells were transfected with the plasmids encoding retromer subunits and treated with DMSO or 1μM XXI for 24 h. Cells were washed with ice cold Dulbecco’s Phosphate-Buffered Saline (DPBS) and lysed in 500 μL of lysis buffer [50 mM HEPES pH7.4, 150 mM NaCl, 1% CHAPSO (Sigma, C3649)] supplemented with 1x HALT protease and phosphatase inhibitor cocktail. The lysate was centrifuged as above and the supernatant was transferred to new tubes. 10% of supernatant was reserved for input samples. The remainder of the supernatant was incubated with 1 μL of anti-PS1 antibody for 3 hours or overnight at 4°C, then incubated with 20 μL of protein G magnetic beads (Thermo Fisher Scientific) for one hour at 4℃. Bound proteins were collected with a magnet, washed four times with lysis buffer, and eluted with 40 μL of 2x Laemmli sample buffer (Bio-Rad) containing 5% 2-mercaptoethanol for 5 min at 100℃, followed by SDS-PAGE and western blot analysis.

### Proximity ligation assay (PLA)

2x10^5^ AAVS and PS1 KO cells or 4x10^5^ HeLa S3 and VPS35/26 KO cells were plated per well in 24-well plates containing glass coverslips and 1 μg of pGFP-DMT1-II or CD8-CIMPR plasmid was transfected into the cells by using Trans-IT HeLaMONSTER reagent or treated with DMSO or 1 μM XXI for 24 h. Cells were fixed, permeabilized, and blocked as described above. Cells were then incubated overnight at 4 ℃ with a pair of mouse and rabbit antibodies recognizing the proteins of interest. PLA was performed with Duolink reagents (Sigma) according to the manufacturer’s instructions. Briefly, cells were incubated in a humidified chamber at 37°C with a pair of PLA antibody probes (mouse and rabbit) for 1 h, with ligation mixture for 45 min, and then with amplification mixture for 3 h, followed by series of washes. Coverslips were mounted with mounting medium containing DAPI. Cellular fluorescence was imaged using the Zeiss LSm980 confocal microscope. Images were processed using a Zeiss Zen software version 3.1 and quantified using Image J Fiji version 1.14.0/1.54f.

### RILP pull down assay

Plasmids expressing GST-RILP (obtained from Christopher Burd, Yale University) in pGEX-KG vector (Addgene, 77103) or pGEX KG expressing GST alone was transformed into *E.coli* strain BL21 (DE3). Bacterial cultures at OD_600_ of 0.6 were induced with 0.4 mM isopropyl β-D-1-thiogalactopyranoside (IPTG) at 30℃ for 6 h. Bacteria were harvested and lysed with B-per lysis buffer (Thermo Fisher Scientific) supplemented with 1x HALT protease inhibitor cocktail (Pierce). GST-RILP or GST proteins were purified by using a pre-equilibrated slurry of glutathione agarose (Thermo Fisher Scientific, 16100) in Buffer I [50 mM Tris pH8.0, 150 mM NaCl, 1 mM MgCl_2_] and washed three times in the same buffer. Purified proteins were eluted from the beads with reduced 20 mM glutathione in Buffer I and dialyzed into HEPES buffer [20 mM HEPES pH7.4, 50 mM NaCl, 5 mM MgCl_2_, 1 mM dithiothreitol (DTT)] using Slide-A-Lyzer dialysis Cassette. Protein amounts were quantified with BSA standards separated by SDS-PAGE followed by Coomassie blue staining.

3x10^6^ AAVS or PS1 KO HeLa cells were plated per dish in 60 mm dishes and transfected with 10 nM of control siRNA (Dharmacon, D-001810-10-05) or TBC1D5 siRNA (Dharmacon, L-020775-01-0005) using Lipofectamine RNAiMAX Transfection Reagent (Thermo Fisher Scientific, 13778100). Cells were then treated with DMSO or 1 μM XXI for 24 h. 48 hpt, cells were washed with ice-cold DPBS and lysed using 300 μL RILP lysis buffer (HEPES buffer containing 0.15% Triton X-100) supplemented with 1x HALT protease and phosphatase inhibitor cocktail. After centrifugation at maximum speed for 20 min at 4℃ in an Eppendorf 5430R centrifuge, the supernatant was incubated with 10 μg of purified GST or GST-RILP proteins at 4℃ for 2 h. 30 μL of mixtures were taken for input samples. Then 40 μL of pre-equilibrated slurry of glutathione agarose was added and the mixtures were further incubated at 4℃ for 3 h. followed by three washes in lysis buffer. Bound proteins were eluted with 40 μL of 2x Laemmli sample buffer containing 5% 2-mercaptoethanol for 5 min at 100℃. Input and eluted samples were separated by SDS-PAGE, and western blot analysis was carried out as described above.

### Statistical analysis

For comparisons of two groups, unpaired *t*-tests were applied. For comparisons of more than two groups, One-way ANOVA with the ordinary ANOVA test were applied. These analyses provide *P-*values for each comparison.

---

**Figure S1.**
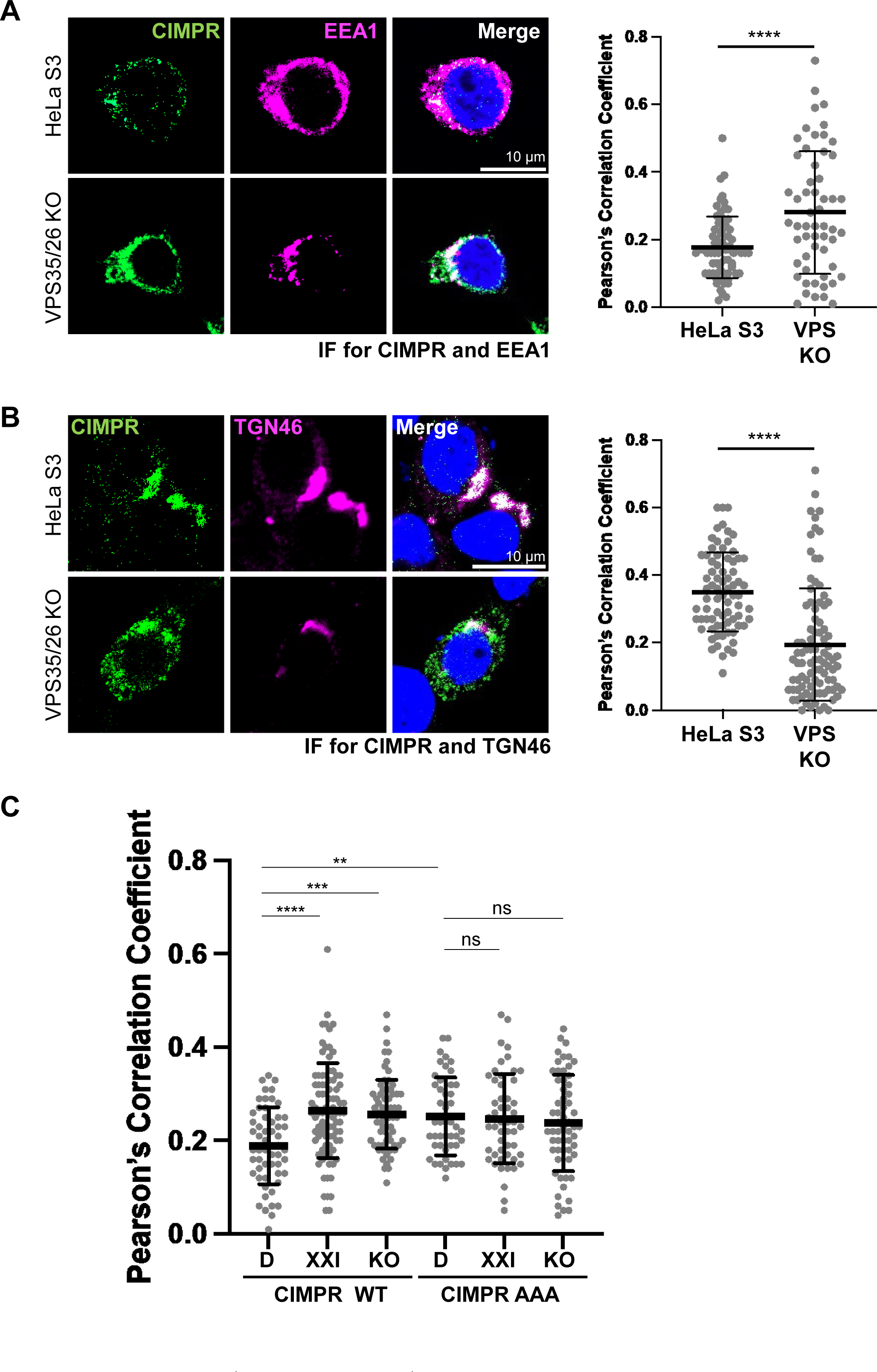
Effect of retromer and γ-secretase on trafficking of wild-type and mutant CIMPR. (**A**) Parental control HeLa S3 cells and VPS35/26 KO HeLa S3 cells (VPS KO) were infected with retrovirus expressing CD8-CIMPR. Cells were fixed 24 h post infection and stained with DAPI and antibodies recognizing CD8 and EEA1. Fluorescent images of single confocal planes are shown: CD8-CIMPR, green; EEA1, magenta; nuclei, blue. Merged images show overlap between CD8-CIMPR and EEA1 pseudocolored white. Graph shows Pearson’s correlation coefficients for colocalization of CD8-CIMPR and EEA1 in cells expressing CD8-CIMPR. Each dot represents an individual cell (n>50), and horizonal line indicates the mean value of the analyzed population in each group. ****, *P*<0.0001. Similar results were obtained in two independent experiments. (**B**) As in (**A**) except cells were stained with antibodies recognizing CD8 and TGN46. Merged images show overlap between CD8-CIMPR and TGN46. (**C**) DMSO-treated AAVS (D), XXI-treated AAVS (XXI), and DMSO-treated PS1 KO HeLa cells (KO) were transfected with a plasmid expressing wild-type CD8-CIMPR (WT) or a AAA mutant CD8-CIMPR (AAA). Cells were fixed 24 hpt and stained with antibodies recognizing CD8 and EEA1. Graph shows Pearson’s correlation coefficients for CD8-CIMPR and EEA1 colocalization as described in (**A**). **, *P*<0.01; ***, *P*<0.001; ****, *P*<0.0001; ns, not significant.

**Figure S2.**
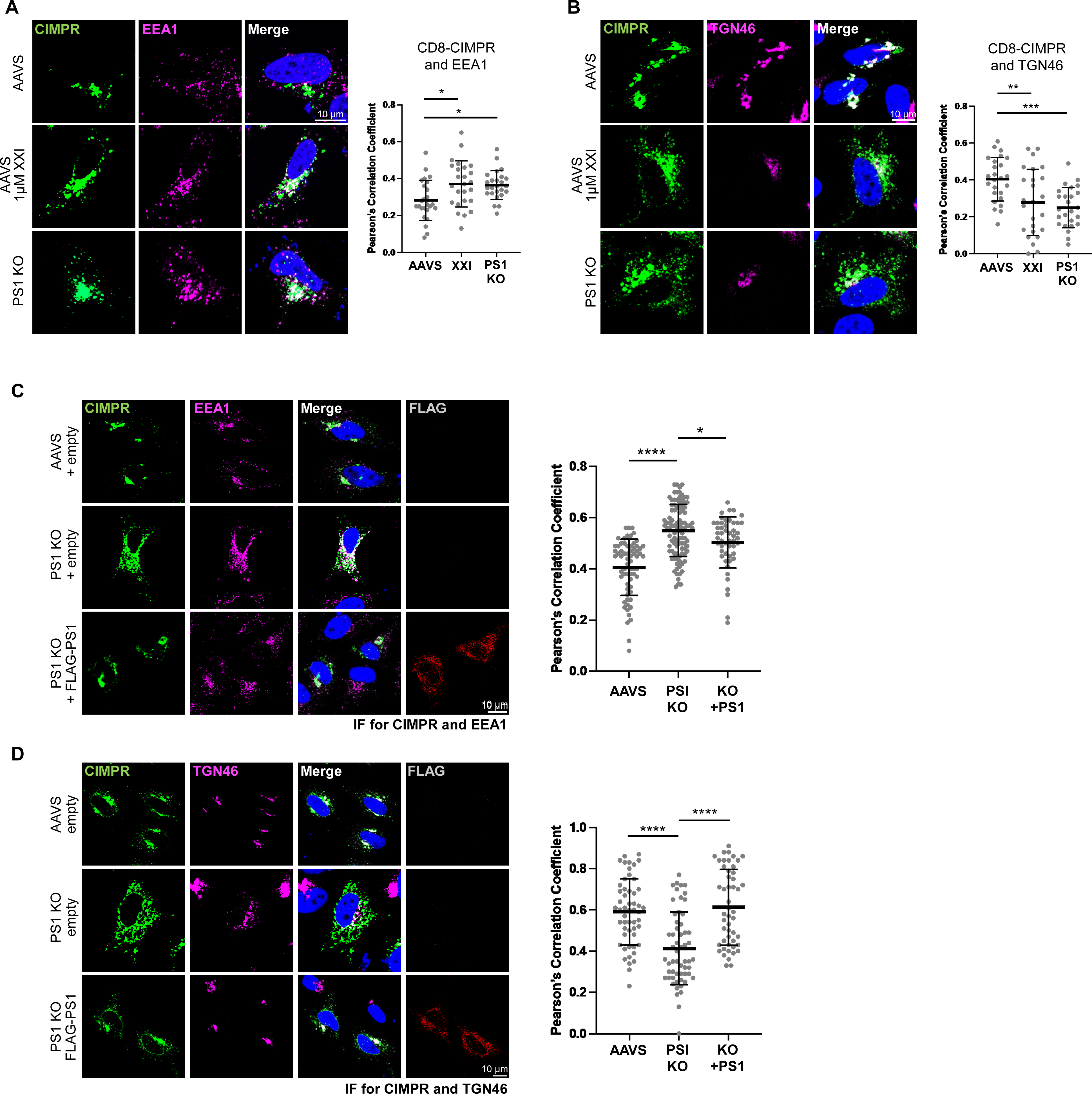
γ-secretase inhibition and PS1 knockout inhibit retrograde trafficking of CIMPR. (**A**) Parental control HeLa cells (AAVS) and PS1 knockout HeLa cells (PS1 KO) were treated with DMSO or 1 μM r XXI for 30 min and then transfected with a plasmid expressing CD8-CIMPR. Cells were fixed 24 hpt and stained with DAPI and an antibody recognizing EEA1. Fluorescence images of single confocal planes are shown: CD8-CIMPR, green; EEA1, magenta; nuclei, blue. Merged images show overlap between CD8-CIMPR and EEA1 pseudocolored white. Graph shows Pearson’s correlation coefficients for colocalization of CD8-CIMPR and EEA1 in cells expressing CD8-CIMPR. Each dot represents an individual cell (n=25), and black horizonal line indicates the mean value of the analyzed population in each group. *, *P*<0.05. Similar results were obtained in three independent experiments. (**B**) As in (**A**) except cells were stained with an antibody recognizing TGN46 instead of EEA1, and merged image shows overlap between CD8-CIMPR and TNG46. **, *P*<0.01; ***, *P*<0.001. (**C**) AAVS and PS1 KO cells were transfected with a plasmid expressing CD8-CIMPR and cotransfected with the empty control plasmid or a plasmid expressing FLAG-PS1. Cells were fixed 24 hpt and stained with DAPI and antibodies recognizing EEA1 and FLAG. Fluorescent images of single confocal planes are shown: CD8-CIMPR, green; EEA1, magenta; FLAG-PS1, red; nuclei, blue. Merged images show overlap between CD8-CIMPR and EEA1 pseudocolored white. Graph shows Pearson’s correlation coefficients for colocalization of CD8-CIMPR and EEA1 in cells expressing CD8-CIMPR (or co-expressing CD8-CIMPR and FLAG-PS1 in the case of cells transfected with plasmid expressing wild-type or mutant PS1). Each dot represents an individual cell (n>50), and horizonal line indicates the mean value of the analyzed population in each group. *, *P*<0.05; ****, *P*<0.0001. Similar results were obtained in two independent experiments. (**D**) As in (**C**) except cells were stained with antibodies recognizing TGN46 and FLAG, and merged images show overlap between CD8-CIMPR and TGN46. ****, *P*<0.0001.

**Figure S3.**
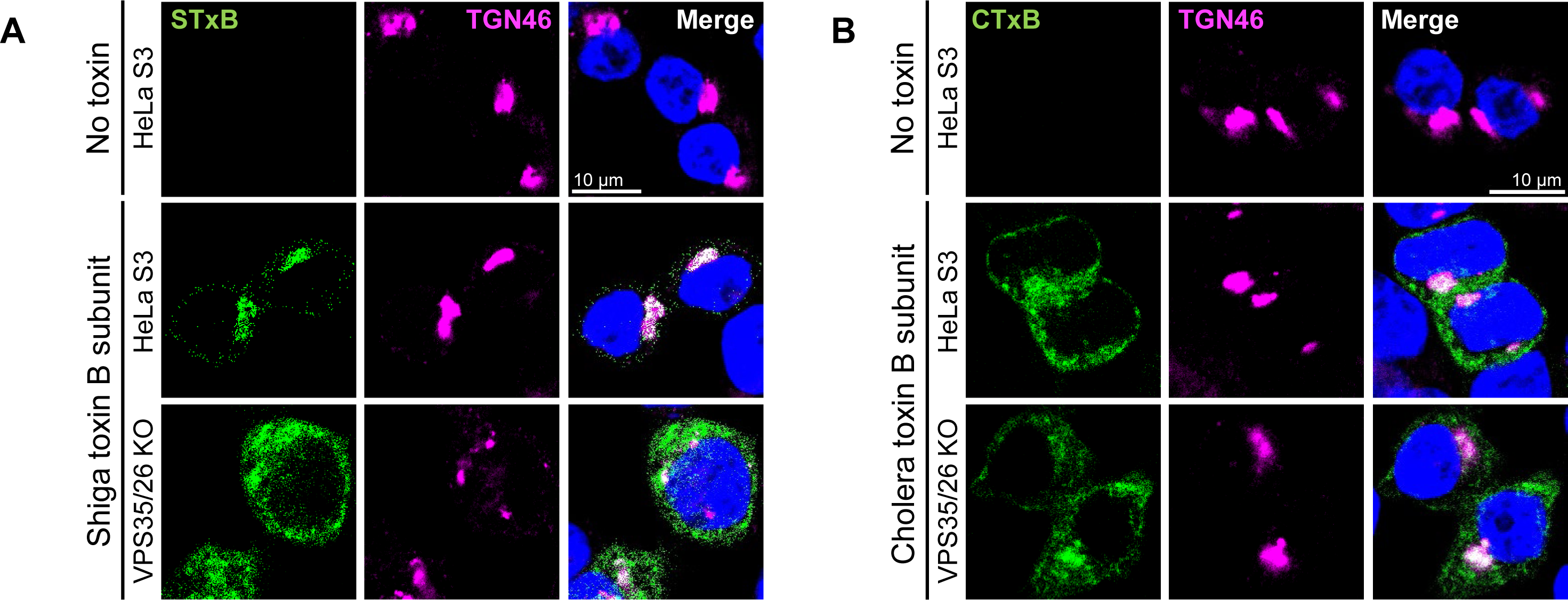
Retromer knockout inhibits retrograde trafficking of shiga toxin but not cholera toxin. (**A**) HeLa S3 cells and VPS26/35 KO cells were incubated with or without 1 μg/ml fluorescent STxB. Cells were fixed 30 min after the treatment and stained with DAPI and an antibody recognizing TGN46. Fluorescent images of single confocal planes are shown: STxB, green; TGN46, magenta; nuclei, blue. Merged image shows overlap between STxB and TGN46 pseudocolored white. Similar results were obtained in two independent experiments. (**B**) As in except cells were incubated with 1 μg/ml fluorescent CTxB. Merged images show overlap between CTxB and TGN46. Quantitation of this experiment is shown in Fig. 2A and B.

**Figure S4.**
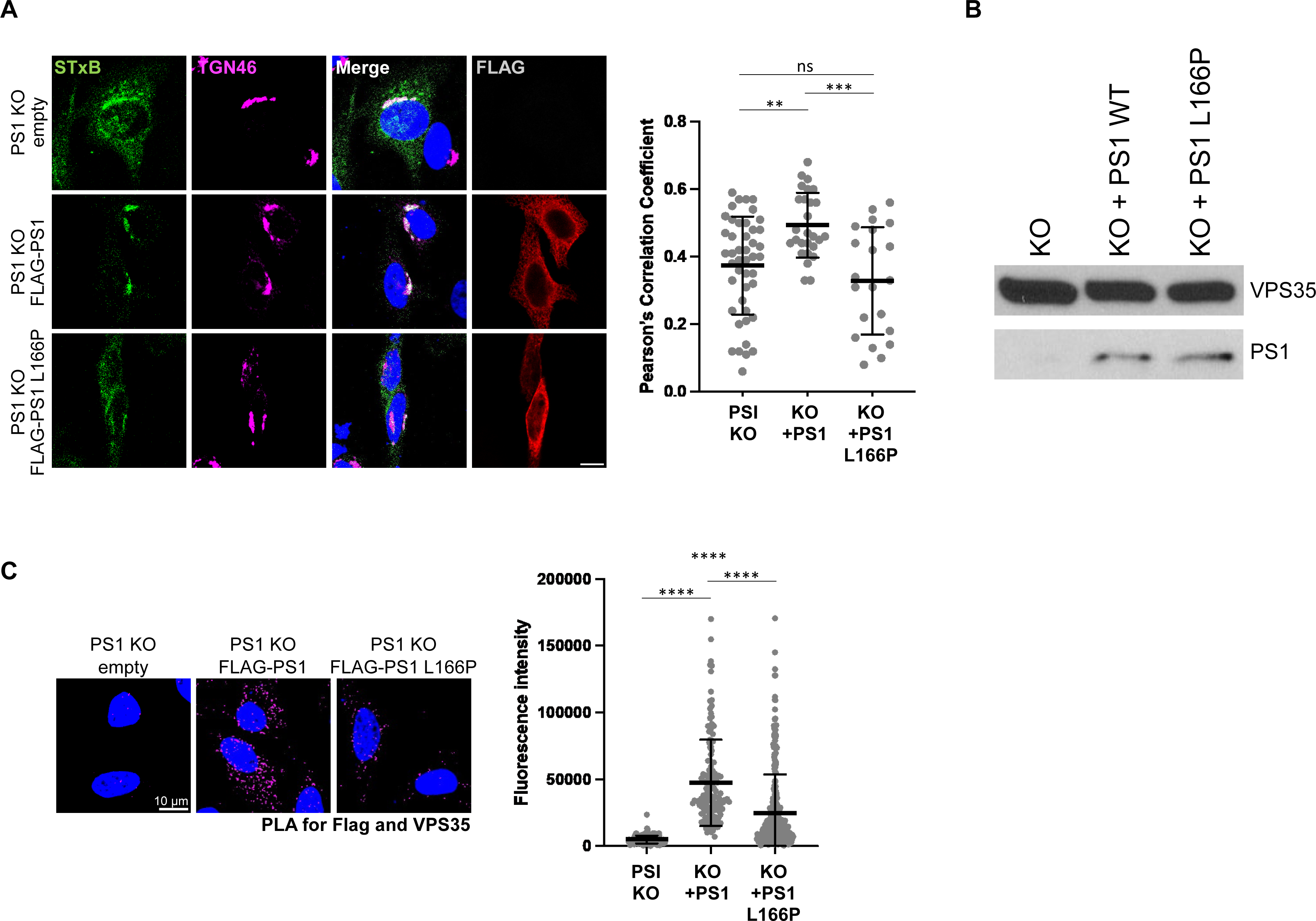
The L166P PS1 mutant does not support trafficking of shiga toxin and is impaired for γ-secretase-retromer interaction. (B) PS1 KO HeLa cells were transfected with the empty control plasmid or a plasmid expressing wild-type FLAG-PS1 or FLAG-PS1 L166P. 24 hpt, cells were incubated with 1 μg/ml fluorescent STxB. Cells were fixed 30 min later and stained with DAPI and antibodies recognizing TGN46 and FLAG. Fluorescent images of single confocal planes are shown: STxB, green; TGN46, magenta; FLAG-PS1, red; nuclei, blue. Merged images show overlap between STxB and TGN46 pseudocolored white. Graph shows Pearson’s correlation coefficients for colocalization of STxB and TGN46 in cells expressing STxB (or co-expressing STxB and FLAG-PS1 or FLAG-PS1 L166P in the case of cells transfected with plasmids expressing wild-type or mutant [KO + PS1 or KO+PS1 L166P]). Each dot represents an individual cell (n>50), and horizonal line indicates the mean value of the analyzed population in each group. ****, *P*<0.0001. Similar results were obtained in two independent experiments. (**B**) Extracts were prepared from PS1 KO HeLa cells transfected with the empty control plasmid or a plasmid expressing FLAG-PS1 or FLAG-PS1 L166P for 24 h. Cell extracts were subjected to western blot analysis using antibodies recognizing endogenous VPS35 and FLAG. (**C**) PS1 KO HeLa cells were transfected with the empty control plasmid or a plasmid expressing FLAG-PS1 or FLAG-PS1 L166P. Cells were fixed 24 hpt. PLA was performed with antibodies recognizing FLAG and VPS35. Images show single confocal planes. PLA signals are magenta; nuclei are stained blue with DAPI. The fluorescence of PLA signals was determined from multiple images. In the graph, each dot represents an individual cell (n>50) and horizonal lines indicate the mean value of the analyzed population in each group. ****, *P*<0.0001. Similar results were obtained in two independent experiments.

